# Placental prostaglandin signaling disrupts barrier integrity and relays an acute inflammatory signal to the fetus

**DOI:** 10.64898/2026.03.11.706681

**Authors:** Hana Horackova, Qiuying Zhao, Shakeela Faulkner, Jennifer Alvarez, Cenk Akiz, Yilin Liu, Josephine Crosthwait, Weiye Dai, Danielle Santoyo, Duke Pham-Chang, Nabeel Bhinderwala, Thea Tagliaferro, W. Dean Wallace, Saloni Walia, Jelena Martinovic, Claire Baldauf, Axel Montagne, Alexandre Bonnin

**Author notes:** These authors contributed equally to this work (Hana Horackova, Qiuying Zhao).

## Abstract

Maternal inflammation during pregnancy is a major risk factor for adverse neurodevelopmental outcomes, yet the mechanisms linking maternal immune activation (MIA) to placenta–fetal brain axis dysfunction remain unclear. Using a poly-(I:C) mouse model, we show that MIA rapidly disrupts placental-blood barrier (PBB) integrity by disrupting pericyte–endothelium coupling within 48 hours, leading to increased placental permeability detected by in vivo MRI. We identify a prostaglandin E2 (PGE2)-dependent pathway involving cyclooxygenase-2 (COX2) and prostaglandin E-synthase (mPGES1) as a key mediator of this effect. Pharmacological inhibition of COX2 prevented MIA-induced structural and functional changes. In vitro explant studies confirmed that PGE2 acts via the EP3 receptor to disrupt pericyte–endothelium interactions, while ex vivo perfusion demonstrated that in addition to its local effects on PBB structure, placenta-derived PGE2 enters the fetal circulation. Finally, similar vascular alterations were observed in human placentas from pregnancies with severe maternal inflammation. These findings reveal a conserved inflammatory mechanism that compromises PBB integrity and may relay signals to the fetal brain, highlighting potential therapeutic targets for neuroprotection during prenatal inflammation.

**One Sentence Summary:** Maternal immune activation during pregnancy disrupts placental vascular integrity through a COX2-dependent prostaglandin E2 pathway, which compromises the placenta–blood barrier and relays inflammatory signals to the fetus, identifying potential therapeutic targets for neuroprotection.

## Introduction

Severe maternal infections during pregnancy—whether viral or bacterial—are epidemiologically associated with later-emerging neurological disorders in offspring^1,2^. In most cases, these effects arise from maternal immune activation (MIA) rather than direct pathogen transmission (with some exceptions, e.g. CMV, Zika virus)^3,4^. A hallmark of MIA is the acute elevation of pro-inflammatory cytokines in maternal circulation^5^. While significant progress has been made in understanding how MIA influences fetal brain development, the role of the placental-blood barrier (PBB)—the primary interface between maternal and fetal compartments—remains underexplored. For MIA to impact fetal neurodevelopment, molecular signals originating in maternal blood must cross two selective barriers: the placenta and the nascent fetal blood-brain barrier (BBB). The placenta not only restricts vertical transmission of pathogens and maternal molecules but also provides essential trophic and hormonal support for brain development^6,7^. Recognition of its role in neurodevelopment has led to the emergence of “neuroplacentology” as a new research field over the past few years^8^.

Using a mouse model, we previously demonstrated that MIA at gestational day (GD) 13 disrupts BBB formation within 48 hours by reducing pericyte coverage of fetal endothelial cells. This disruption caused chronic BBB breakdown, hyperpermeability, parenchymal inflammation, and behavioral alterations persisting into adulthood^9^. We also observed increased proliferation and perivascular localization of cyclooxygenase-2 (COX2; *Ptgs2*)-expressing microglia. Genetic deletion of Ptgs2 or pharmacological inhibition of COX2 prevented these changes, implicating prenatal COX2 activation as a key mediator of MIA-induced neurodevelopmental abnormalities^9^. Although the upstream signals driving COX2 activation in the fetal brain remain unclear prostaglandins themselves (e.g. PGE2)^10^ and/or cytokines (e.g. IL6)^11^ are strong candidates.

The PBB shares structural features with the BBB, including pericyte-like cells covering fetal endothelial cells, but also includes additional layers of syncytiotrophoblasts (SCTs) that separate maternal and fetal blood (Fig. 1A). COX2 is expressed in the placenta, where it regulates vascular function and plays a critical role in parturition^12^. These similarities led us to hypothesize that MIA disrupts placental vascular integrity through COX2-dependent mechanisms. To test this, we employed a poly(I:C)-induced MIA mouse model and combined live MRI, histological analysis, functional *ex vivo* perfusions and *in vitro* studies. Our findings reveal that MIA activates the COX2 pathway via PGE2 signaling through the EP3 receptor (*Ptger3*) in pericyte-like cells surrounding fetal endothelium, leading to PBB disruption. Similar vascular changes were observed in a small cohort of human pregnancies complicated by severe MIA. Importantly, *ex vivo* placental perfusion experiments demonstrated that placenta-derived PGE2 enters the fetal circulation, identifying a potential molecular link between placental inflammation and fetal BBB dysfunction.

**Figure 1.**
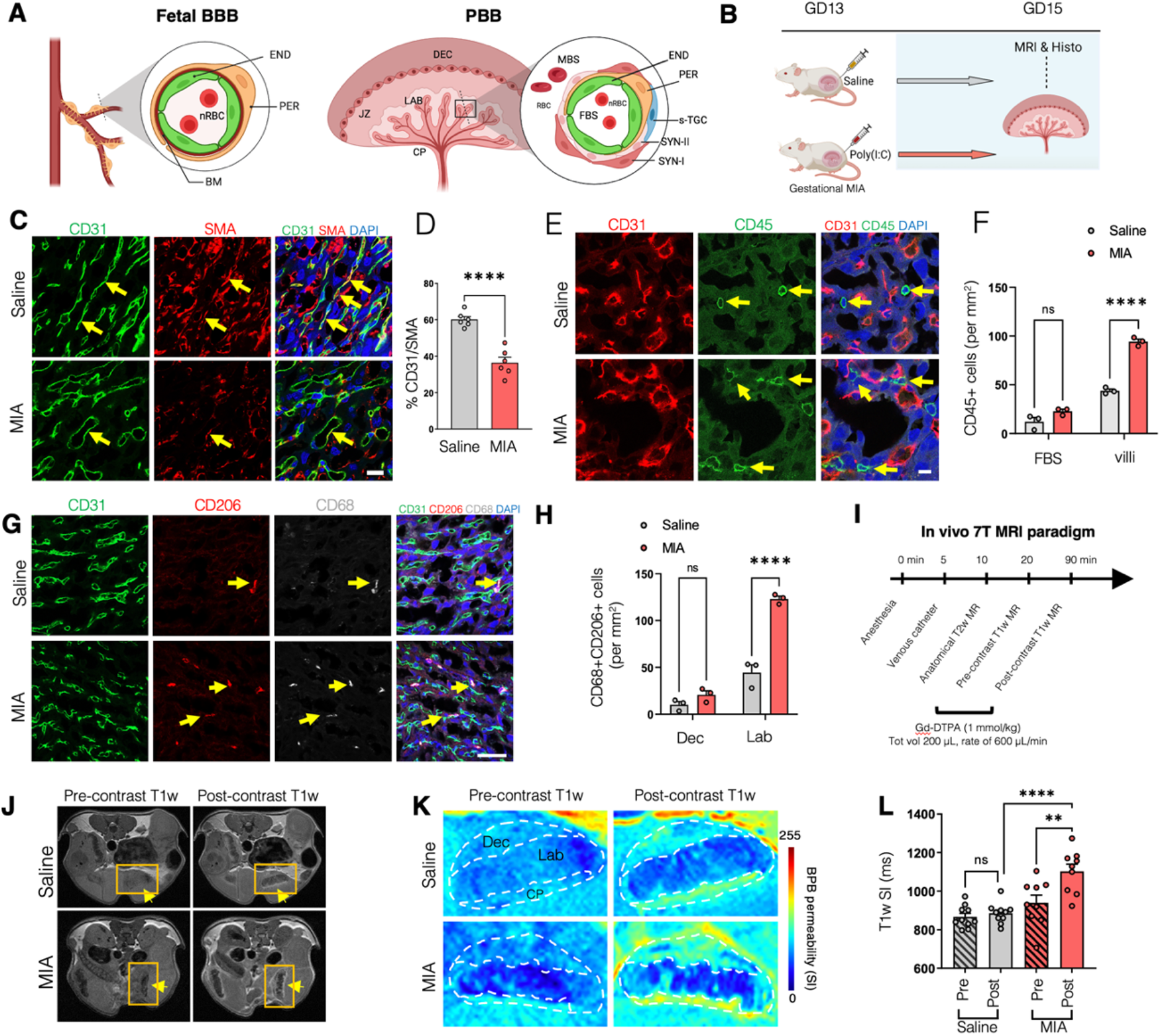
MIA induces PBB vascular breakdown. (**A**) Comparison of mouse fetal BBB and PBB structures (END, endothelium; PER, pericytes; BM, basement membrane; n/RBC, nucleated/red blood cells; M/FBS, maternal/fetal blood space; s-TGC, sinusoidal trophoblastic giant cell; SYN I/II, syncytiotrophoblast I/II; Dec, decidua; JZ, junctional zone; Lab, labyrinth; CP, chorionic plate). (**B**) Pregnant CD-1 mice received saline or poly(I:C) on GD13; effects were measured at GD15. (**C**) Colocalization (yellow arrows) of pericyte-like cell marker SMA (red) and endothelial cells marker CD31 (green) in GD15 saline and MIA placentas. Scale bar, 20µm. (**D**) Quantification of pericyte coverage of endothelial cells (SMA-CD31 signal colocalization) in GD15 placenta. n=6 dams/group, one placenta per dam. (**E-F**) Representative images and quantification of CD45+ macrophages (arrows) in GD15 labyrinth villi and FBS. Scale bar, 10µm. (**G-H**) Representative images and quantification of resident (CD206+, arrows) activated macrophages (CD68+, grey) in GD15 placental labyrinth and decidua. Scale bar, 50µm. n=3 dams/group, one placenta per dam. (**I-J**) MRI timeline and example of T1w pre- and post-contrast images in saline and MIA mice (arrows show placentas). (**K**) Heatmap of T1w pre- and post-contrast images demonstrates increased contrast signal in the MIA placenta. (**L**) Quantification of T1w SI pre- and post-contrast in saline (n=10 dams) and MIA (n=9 dams), 1-3 placentas analyzed per dam. Data represent mean ± SEM and evaluated by one-way ANOVA with Tukey’s post-hoc test (**L**), two-way ANOVA with Bonferroni’s correction for multiple comparisons (**F, H**) or two-tailed t test (**D**); **p < 0.01, ****p < 0.0001.

## Results

### Gestational MIA disrupts placenta-blood barrier vascular structure and function

Maternal immune activation (MIA) is known to impair pericyte–endothelium coupling in the fetal blood–brain barrier (BBB^9^). Because the fetal BBB and placenta–blood barrier (PBB) share similar vascular architecture (Fig. 1A), we investigated whether MIA also disrupts this coupling in the placenta at gestational day (GD) 15. MIA was induced by poly(I:C) injection at GD13 (Fig. 1B).

Histological analysis revealed a significant reduction in pericyte-like cell coverage of endothelial cells [a-smooth muscle actin SMA/ACTA2-positive pericytes^13^ and CD31-positive endothelium) 48 hours after poly(I:C) injection compared to saline controls (Fig. 1C–D). Fetal sex did not significantly affect pericyte–endothelium coupling (P = 0.74, F(1,8) = 0.11; n = 3 per sex; two-way ANOVA with Bonferroni correction). Additionally, MIA increased the number of CD45-positive macrophages near fetal vessels in the placental labyrinth, but not in maternal blood spaces or decidua (Fig. 1E–F). Further characterization identified these cells as activated placental resident macrophages (CD206+/CD68+, Hoffbauer cells^14^; Fig. 1G–H). These findings indicate that MIA disrupts pericyte–endothelium interactions and triggers local inflammatory responses in the placenta.

To assess whether reduced pericyte coverage alters PBB permeability, we performed live MRI using a 940 Da tracer (Gd-DTPA) to measure diffusion from maternal to fetal blood compartments over 90 minutes. High-resolution abdominal T1-weighted imaging quantified signal intensities in the labyrinth and chorionic plate before and after intravenous Gd-DTPA injection (Fig. 1I–K). At GD15, poly(I:C)-treated placentas exhibited increased Gd-DTPA diffusion 70 minutes post-injection compared to saline controls (Fig. 1L), indicating that MIA-induced vascular disruption compromises placental barrier integrity.

### MIA effect depends on prostaglandin pathway activation

Prostaglandin signaling mediates vascular breakdown in the fetal BBB^9^. To test whether PBB disruption also depends on COX2 activity, dams received celecoxib (a COX2 inhibitor) or vehicle (DMSO) 24 hours after poly(I:C) or saline injection (Fig. 2A). Celecoxib treatment restored pericyte coverage to control levels (Fig. 2B–C), prevented macrophage activation (Fig. 2D–E), and normalized Gd-DTPA diffusion in vivo (Fig. 2F). These results confirm that COX2 activation is required for MIA-induced PBB disruption.

**Figure 2.**
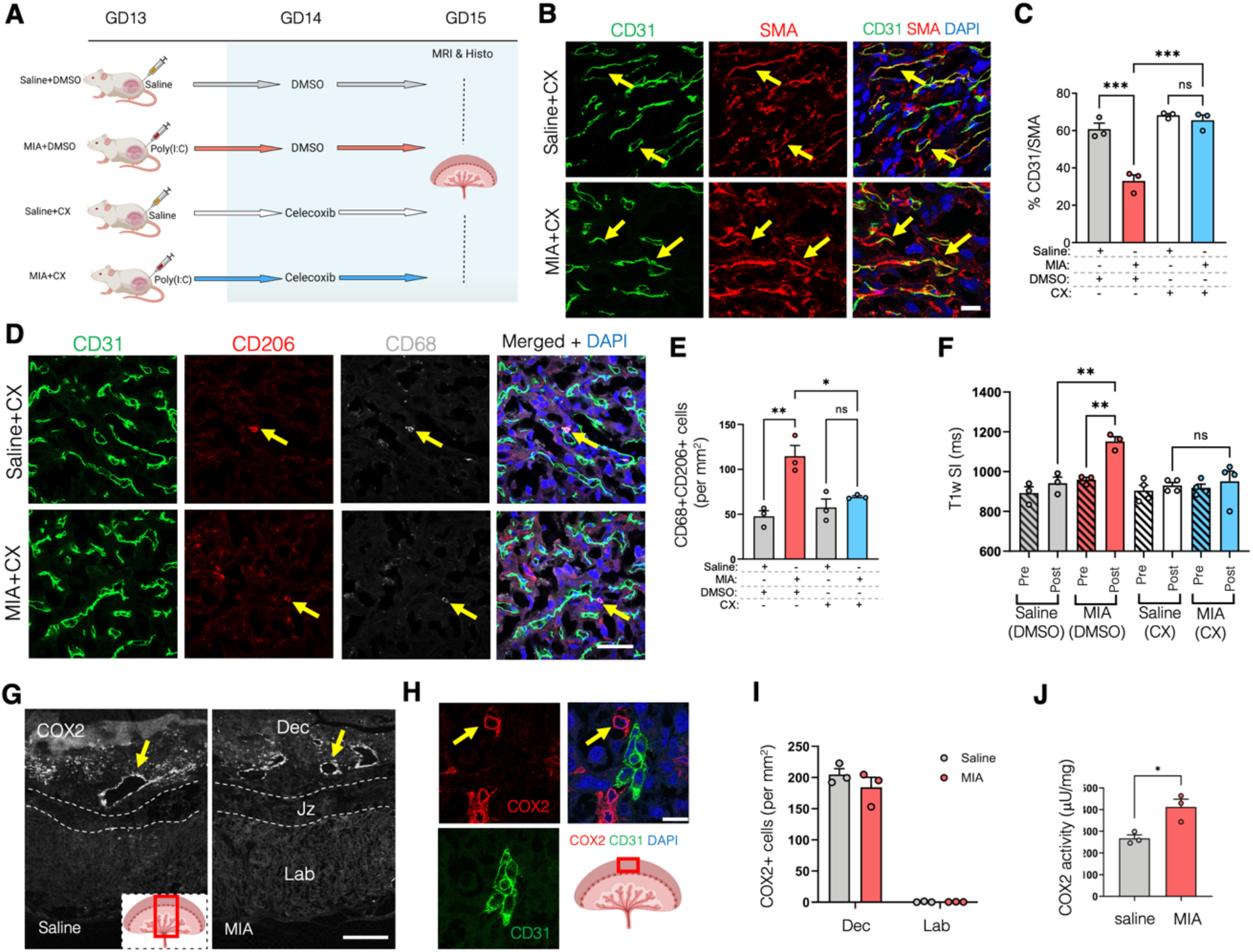
COX2 mediates MIA effects on PBB function. (**A**) Pregnant CD-1 mice were intraperitoneally (i.p.) injected once on gestational day GD 13 with saline or poly(I:C); 13.8 mg/kg and subsequently (i.p.) with DMSO or the COX-2 inhibitor celecoxib at GD14. Samples were collected at GD15 for MRI and histology. Blocking of COX2 activity prevented MIA effect on SMA-CD31 signal colocalization, macrophage activation and PBB permeability to Gd-DTPA. (**B**) Representative images of endothelial (CD31+, green) and pericyte-like (SMA+, red) cells colocalization (yellow arrows) in GD15 placenta of mice treated with celecoxib (CX) 24h after saline or poly(I:C) (MIA) injection, scale bar, 20µm. (**C**) Quantification of SMA-CD31 signal overlap in placenta at GD15. (**D**) Representative images of macrophage markers (CD206+ red, CD68+ grey) in GD15 labyrinth of placenta. (**E**) Quantification of CD206+ and CD68+ double positive cells in the labyrinth at GD15, n=3 dams/group, one placenta per dam. (**F**) Quantification of T1w SI pre- and post-contrast at GD15. MIA placentas without CX treatment show increased permeability vs treated MIA and controls. (**G)** COX2+ cells located around CD31+ maternal vessels (arrows) in untreated saline or MIA decidua (scale bar, 500µm), and higher magnification immunofluorescence image in **H** (scale bar, 40 µm). (**I**) Quantification of COX2+ cells in the decidua (Dec) and labyrinth (Lab) of GD15 placentas from saline or MIA treated dams. (**J**) COX2 enzymatic activity in placenta 28 h after GD13 saline or poly(I:C) injection, n=3 dams/group, one placenta per dam. Data are presented as mean ± SEM (\**p* < 0.05, \*\**p* < 0.01, and \*\*\**p* < 0.001 and evaluated by one-way ANOVA with Tukey’s post-hoc comparison test (**C, E, F**) or two-way ANOVA with Bonferroni’s correction for multiple comparisons (**J**).

In the placenta, COX2 was absent in resident macrophages but expressed in decidual stromal cells surrounding maternal arteries (Fig. 2G–H). COX2 enzymatic activity increased 28 hours after poly(I:C) injection without changes in COX2-positive cell numbers (Fig. 2I–J). This suggests that prostaglandins produced in the decidua may reach the labyrinth via maternal blood flow.

### Mapping Prostaglandin Pathway Components

We next examined expression of microsomal prostaglandin synthase-1 (mPGES1), the terminal enzyme in PGE2 synthesis (Fig. 3A). Western blot analysis revealed highest mPGES1 expression in the decidua, with significant downregulation at GD15 following MIA (Fig. 3B–C). Immunohistochemistry localized mPGES1 to syncytiotrophoblast (SCT) layers within the labyrinth, positioned between MCT1-positive SCT I/II and CD31-positive endothelium (Fig. 3D). This suggests SCTs convert COX2-derived PGH2 to PGE2, which may influence pericyte–endothelium coupling and enter the fetal circulation.

**Figure 3.**
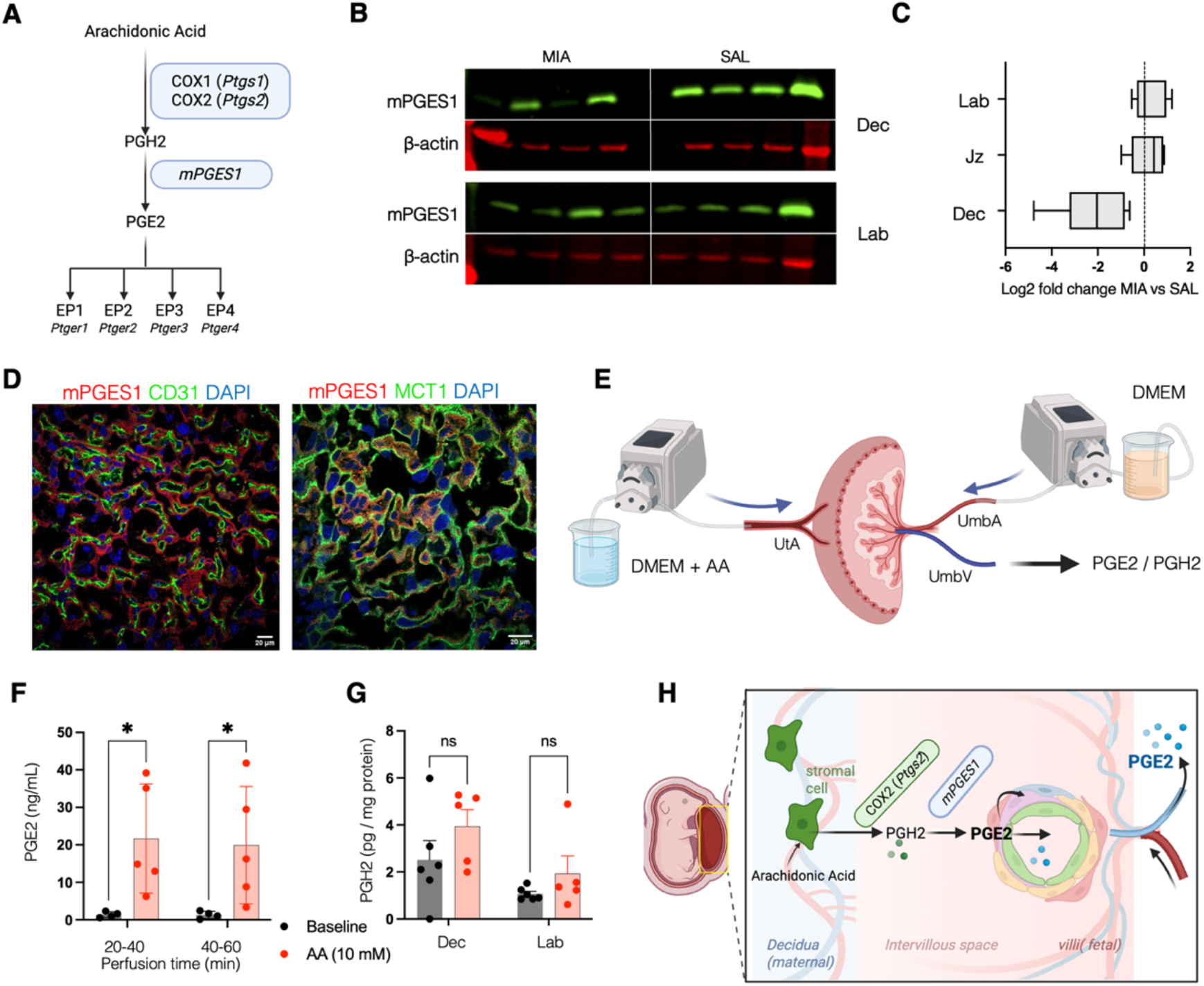
COX2 and mPGES1 function in the PBB. (**A**) Arachidonic acid (AA) to PGE2 enzymatic pathway. (**B**) Representative western blots of mPGES1 protein expression in placental layers in saline (SAL) and MIA homogenates. (**C**) Log2 fold decrease of relative mPGES1 protein expression in MIA vs saline Dec, JZ and Lab. Protein expression was normalized to β-actin. (**D**) mPGES1 (red) immunostaining localized between MCT1+ (Syncytiotrophoblast I marker) and CD31+ endothelial cells (green). (**E**) Diagram of *ex vivo* placenta perfusion with AA. (**F**) PGE2 concentration in fetal perfusates in baseline and AA perfusions. (**G**) PGH2 concentration in Dec and Lab of control placentas after baseline and AA (10 mM) perfusion. (**H**) Illustration of AA to PGE2 pathway in MIA placenta. Log2 fold change of mPGES1 presented as ± SEM (**p=0.0031) and evaluated by one-sample t and Wilcoxon test (**C**). PGE2 and PGH2 concentrations presented as ± SEM and evaluated by two-way ANOVA (\**p* < 0.05).

To test this, we perfused GD14 placentas *ex vivo* via the uterine artery^15^ with arachidonic acid (AA, 2.5 mg/mL) and measured prostaglandins in the umbilical vein (Fig. 3E). PGH2 was undetectable in fetal perfusate, but AA infusion significantly increased PGE2 levels at 20–40 and 40–60 minutes (Fig. 3F). Post-perfusion tissue analysis showed a trend in PGH2 accumulation in decidua and labyrinth (Fig. 3G). These findings reveal a pathway where AA is converted to PGH2 by COX2 in the decidua, then to PGE2 by mPGES1 in SCTs, with PGE2 released into fetal circulation (Fig. 3H).

### EP3 Receptor Mediates Vascular Disruption

Besides its release into the fetal circulation, SCT-derived PGE2 may act upon the proximal pericyte-endothelium complex via a variety of prostaglandin receptors (EP1-4; *Ptger1-4*, Fig. 3A). Immunohistochemistry showed EP3 receptor localization in the labyrinth, primarily in pericyte-like cells (Fig. 4A), positioning it to mediate PGE2 effects. EP3 expression decreased significantly 48 hours after MIA (Fig. 4B). GD14 placental labyrinth explants incubated with PGE2 (100 pg/mL) or EP3 agonist sulprostone (10 μM) for 48 hours exhibited reduced pericyte– endothelium coverage (Fig. 4C–E). PGH2 incubation led to rapid PGE2 synthesis (524 ng/mL released in culture medium within 3 hours) and decreased SMA/CD31 coverage after 48 hours (Fig. 4F–G). These results confirm that local PGE2 synthesis disrupts pericyte–endothelium interactions via EP3 signaling (Fig. 4H).

**Figure 4.**
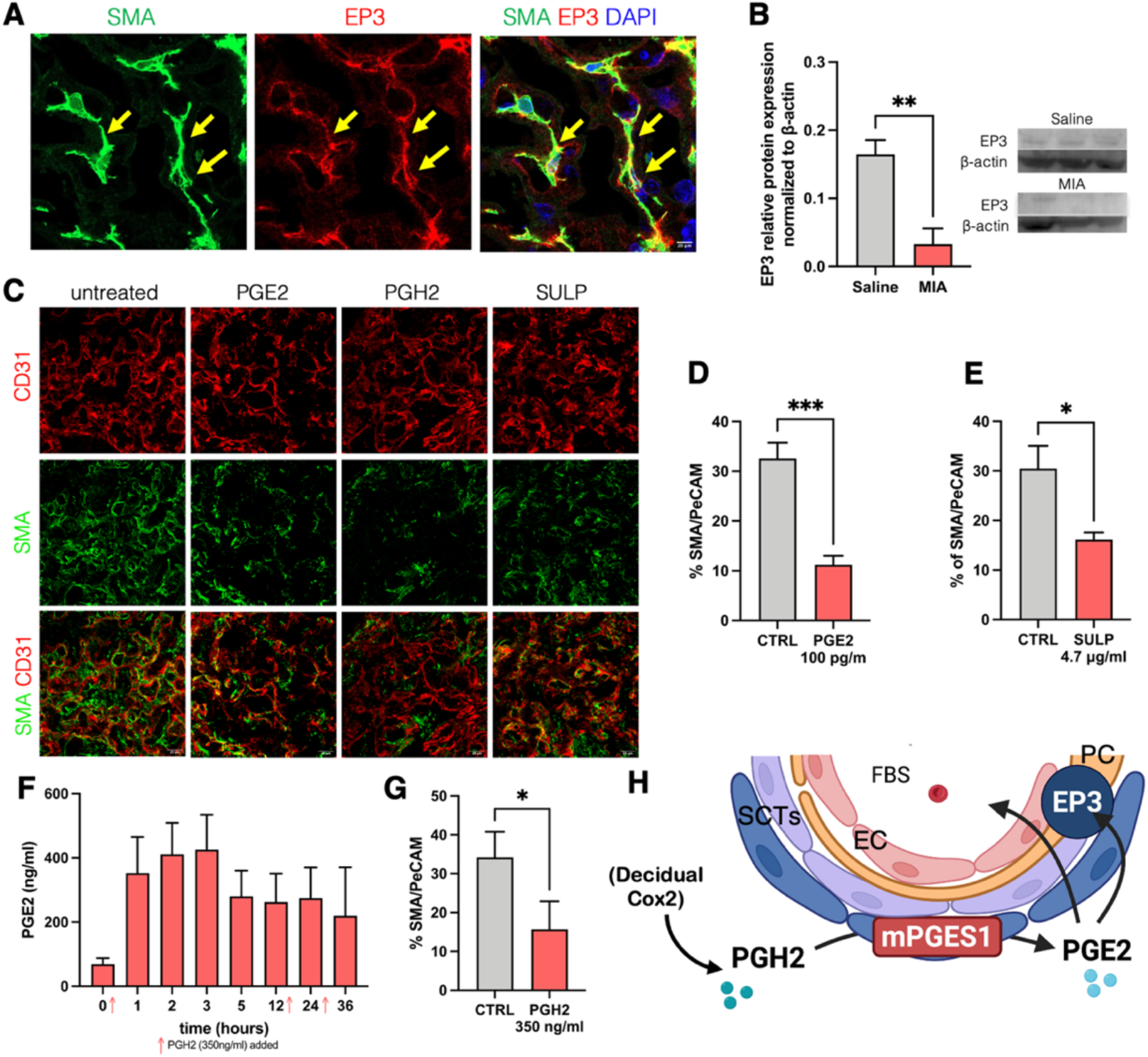
PBB disruption is mediated by EP3 receptor activation. (**A**) Colocalization (arrows) of EP3 receptor (red) with pericyte-like cell marker SMA (green) and nuclear stain DAPI (blue) in labyrinth of control placentas (Scale bar, 20 µm). (**B**) Quantification of EP3 receptor protein expression in the placental labyrinth normalized to β-actin in saline versus MIA groups (N=3/group). (**C**) Representative immunoblots for EP3 and β-actin are shown. Representative images showing SMA-CD31 staining overlap in mouse placenta labyrinth explant cultures: Untreated, treated with PGE2 (100 pg/mL), PGH2 (350 ng/mL), or EP3 receptor agonist sulprostone (SULP, 4.7 µg/mL) for 48 hours. Scale bar, 20 µm. (**D-G**) Quantification of pericyte-like coverage of endothelial cells in cultures treated with PGE2 (**D**), sulprostone (**E**) and PGH2 (**G**). (**F**) PGE2 concentration in media from explants treated with PGH2 (350 ng/mL). Red arrows indicate medium change with addition of fresh PGH2. N=3 independent cultures per condition. (**H**) Schematics of the proposed intra-placental prostaglandin signaling cascade mediating the loss of pericyte coverage in MIA placenta. FBS = fetal blood space, SCTs = syncytiotrophoblast. Data presented as mean ± SEM and evaluated by unpaired two-tailed t-test; \**p* < 0.05, \*\**p* < 0.01, and \*\*\**p* < 0.001.

### Maternal Infection Disrupts Placental Vascular Structure in Humans

To assess clinical relevance, we analyzed placentas from mothers with chorioamnionitis or HIV antepartum infection (n = 6), PCR-positive asymptomatic COVID-19 (n = 8), and controls (n = 5). Multiplex immunofluorescence revealed significantly reduced pericyte–endothelium coverage and increased CD68+ macrophages in placentas from HIV and chorioamnionitis cases, but not in asymptomatic COVID-19 or controls (Fig. 5A–C). These findings suggest that severe maternal infections, but not mild viral exposure, can disrupt PBB integrity in humans.

**Figure 5.**
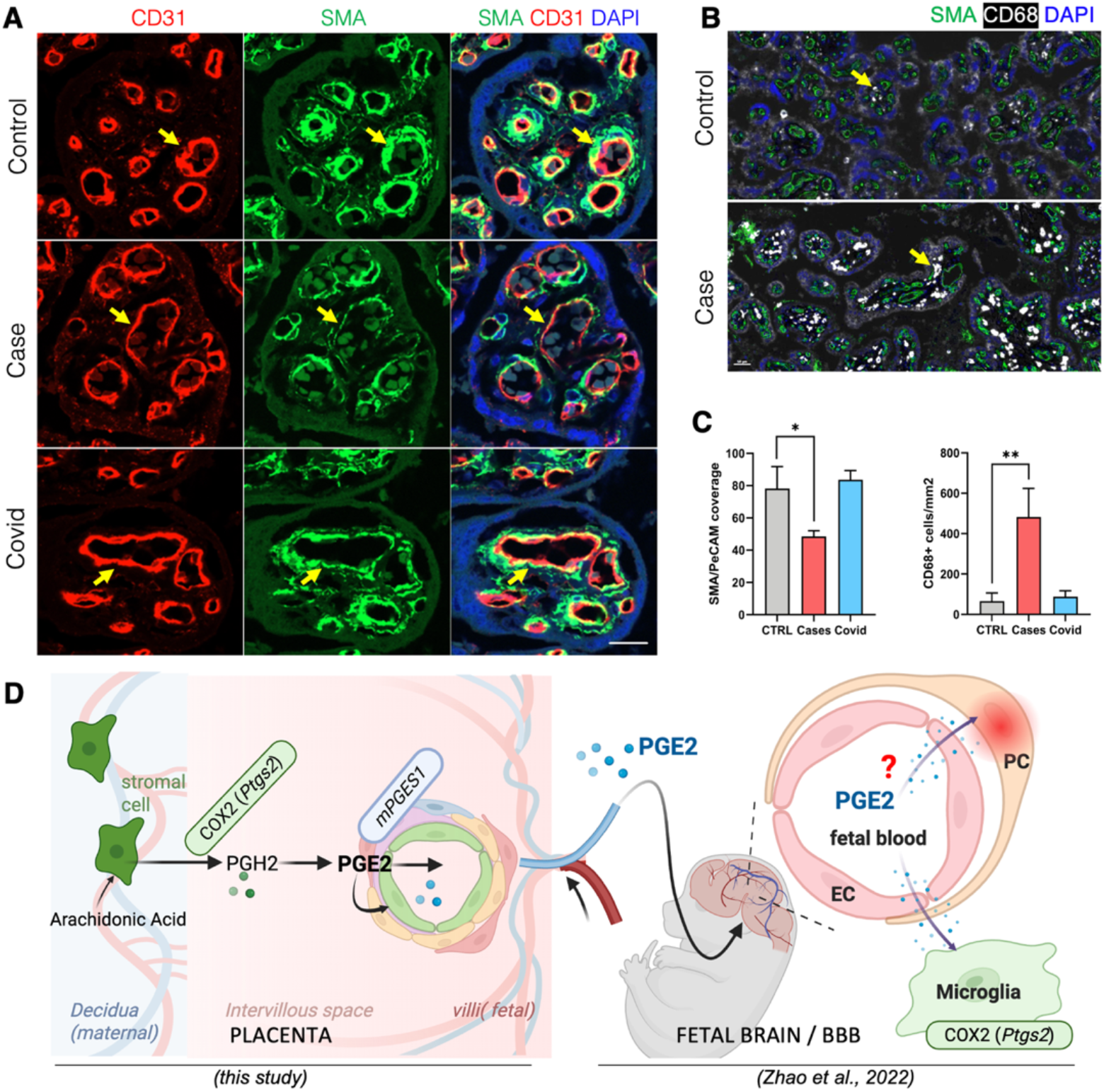
PBB disruption and increased macrophages density in human placenta with inflammation but not with asymptomatic COVID-19. (**A**) Representative images of overlap (arrows) between pericyte-like cell marker SMA (green) and endothelial cell marker CD31 (red) in placentas from human healthy controls, cases of antepartum infection (Case) and asymptomatic COVID 19 (Covid). Scale bar, 20µm. (**B**) CD68+ macrophages (arrow) in fetal villi in a representative control and case (inflammation) placenta. (**C**) Quantification of SMA-CD31 signal overlap and of CD68+ macrophages in human placenta villi. Control (n=5), Cases (n=6), COVID 19 (n=8). (**D**) Illustration of the proposed physiological pathway by which MIA activation of the COX2 pathway in the placenta might not only alter vascular structure in the PBB but also in the fetal BBB. Data are presented as median + IQR, Kruskal-Wallis test with Dunn’s correction for multiple comparison, \**p* < 0.05, \*\**p* < 0.01

## Discussion

### Summary of Main Findings

Our study identifies a prostaglandin-dependent mechanism by which maternal immune activation (MIA) compromises placental barrier function. Within 48 hours of MIA induction in mice, we observed disrupted interactions between pericyte-like cells and endothelial cells in the placental blood barrier (PBB), leading to increased placental permeability *in vivo*. Pharmacological inhibition of COX2—the primary enzyme for prostaglandin E2 (PGE2) synthesis—prevented these effects.

Localization studies revealed that mPGES1 (a secondary PGE2-synthesizing enzyme) and the PGE2 receptor EP3 are concentrated in the labyrinth region. *In vitro* experiments using labyrinthine explants confirmed that pharmacological activation of this pathway rapidly disrupts PBB vascular structure within 24 hours. Similar vascular alterations were observed in human placentas exposed to antepartum infections.

Inflammation is known to impair vascular barriers such as the blood-retina barrier^16^ and the blood-brain barrier (BBB) in both pre-^9^ and postnatal stages^17,18,19^. Evidence suggests that maternal inflammation affects endothelial tight junctions in the PBB, increasing permeability through pro-inflammatory cytokines. Intrauterine infections (e.g., TORCH^20^, CMV^21^), and systemic inflammatory diseases alter placental adhesion molecules and vascular morphology, often reducing pericyte coverage^22^. Murine models of maternal influenza infection have shown placental thrombosis and structural disorganization^23^, with leakage of large molecular tracers into the fetal compartment, including the brain^24^.

### Mechanistic Insights

Our results using live MRI and histological data revealed a rapid disruption of PBB function by viral-like induced MIA, evidenced by increased diffusion of maternal Gd-DTPA tracer and decreased coverage of endothelial cells by pericyte-like cells in the placenta labyrinth. Pericytes are involved in maintaining vascular stability and permeability in the BBB, but the role of their homologs (pericyte-like or Rouget cells) in placental function is less clear. Analogous to the fetal BBB in terms of structural and functional consequences of MIA, our data suggest that pericyte-like cell interactions with fetal endothelium are important for maintaining PBB integrity.

Live MRI and histological analyses demonstrated that viral-like MIA rapidly disrupts PBB integrity, as indicated by increased maternal Gd-DTPA tracer diffusion and reduced pericyte-like coverage of endothelial cells. Pericytes maintain vascular stability in the BBB^25^, but their placental counterparts (Rouget cells^26^) are less understood. Consistent with recent studies suggesting a role for pericyte-like cells in placental vascular integrity^27,28^, our findings suggest these cells are critical for PBB homeostasis.

Furthermore, we observed that COX2 pathway activation is essential for MIA-induced PBB disruption. Unlike the fetal brain where COX2 enzyme is expressed by microglia^9^, placental COX2 is localized mainly in decidual stromal cells near maternal arteries. Hofbauer cells (placental macrophages^29^) increased in number after MIA but did not express COX2, indicating that in contrast to an *in vitro* model^30^, they may respond to prostaglandin signals rather than produce them. This response may include vascular damage repair via their recruitment to fetal blood vessels in response to PGE2 released from SCTs (see below). Similarly, in the brain, an increase in perivascular microglia (brain resident macrophages) was observed around the fetal BBB after MIA^9^. An elegant study of adult BBB demonstrated that in response to inflammation, vessel-associated microglia initially attempt to maintain BBB integrity via expression of the tight-junction protein Claudin-5 and make physical contact with endothelial cells^31^. Our observations in the PBB could reflect an analogous response for Hofbauer cells, with involvement in a rapid repair mechanism upon loss of pericyte-like cell coverage of endothelial cells with the aim of preserving barrier function. This would be consistent with Hofbauer cells’ role in protecting the fetus against the effects of inflammation during pregnancy^32,33^.

### Proposed Mechanism

Our data support the following sequence (illustrated Fig. 5D): 1) MIA increases COX2 activity in the decidua, producing PGH2. 2) PGH2 enters the maternal blood compartment and is converted to PGE2 by mPGES1 in syncytiotrophoblasts. 3) PGE2 activates EP3 receptors on pericyte-like cells, disrupting their interaction with fetal endothelial cells, and 4) some placental PGE2 enters the fetal circulation, potentially signaling to the fetal BBB and contributing to neurodevelopmental effects. This mechanism was confirmed through *ex vivo* perfusion and *in vitro* pharmacological experiments. Notably, COX2 activity increased without changes in COX2+ cell numbers, suggesting regulation occurs via substrate availability or post-translational mechanisms^34^ rather than protein expression.

### Human Relevance

Analysis of a limited number of human placentas from pregnancies complicated by severe inflammation (HIV, chorioamnionitis) revealed increased CD68+ macrophages and reduced pericyte-like coverage, mirroring murine findings. Placentas from asymptomatic SARS-CoV-2–positive women showed no structural changes, indicating that maternal inflammatory response—not infection alone^35,36^—activates the COX2 pathway.

### Physiological and Therapeutic Implications

There are several important physiological implications of our findings. First, the demonstration that placentally derived PGE2 is released into the fetal blood stream suggests that during MIA induction, PGE2 is the initial signal relaying the inflammatory trigger from the PBB to the fetal BBB^9^. This is consistent with the rapid post-MIA protective effects of COX2 inhibition (via Celecoxib) on both PBB and BBB structure and function. Though elevated maternal serum cytokines are generally assumed to provide inflammatory relay signals, there is little evidence of actual transplacental transfer of maternal cytokines to the fetal brain^37,38^. Instead, cytokines such as IL-6, which are dramatically increased in the maternal serum within 6h of MIA induction in our model^9^ may provide an IL-6 receptor mediated^39^ trigger for the activation of the placental prostaglandin pathway demonstrated here. Various immunological systems have demonstrated cytokine mediated prostaglandin pathway activation^40^. However, to rapidly relay inflammatory signals across the placenta to the fetus, placenta-derived prostaglandin are ideally suited, because unlike multi-kDa cytokine proteins, PGE2 is a small lipophilic molecule that that easily crosses cellular membranes^41^.

Second, EP3 receptor antagonism may offer a targeted therapeutic strategy to protect fetal development, avoiding risks associated with broad COX2 inhibition (e.g., premature ductus arteriosus closure^42^). Future studies to elucidate the effect of EP3 antagonism on fetal ductal tissue and circulating levels of fetal PGE2 would further inform the therapeutic potential of EP3 as a target for MIA. Lastly, follow up studies examining inhibition of the COX2 pathway at varying gestational ages and additional post-MIA timepoints with attention to impact on the fetal BBB are warranted.

### Conclusion

Maternal immune activation compromises placental vascular integrity through a COX2-dependent prostaglandin pathway that disrupts pericyte-endothelial interactions. Targeting EP3 signaling or placental prostaglandin release represents a promising approach to preserve placental function and protect fetal brain development.

## Acknowledgements

Supported by NIH (5R01NS126981 to A.B.), BrightFocus Foundation (#A2019279S to A.B.), Cure Alzheimer’s Foundation (to A.B.), and CIRM (EDUC4-12802 to H.H.). We thank Dr. Juli Wu for scientific editing. This work was supported by the Cellular Imaging Core of the Saban Research Institute at Children’s Hospital Los Angeles (CHLA). The content is solely the responsibility of the authors and does not necessarily represent the official views of the NIH.

## Author Contributions

H.H., Q.Z. and A.B. designed the experimental studies. H.H., Q.Z. conducted the experiments. J.A. conducted the ex vivo perfusion experiments. J.C., C.A., Y.L., W.D., D.S., D.P.C. performed immunohistochemical and blot analyses. S.F., N.B., T.T., W.D.W., S.W., J.M. and C.B. provided clinical samples and performed human placenta analyses. H.H., Q.Z., A.M. and A.B. analyzed the data and reviewed the results. H.H., Q.Z., C.B. and A.B. wrote and edited the manuscript.

## Material and Methods

### Animals

CD-1 timed pregnant mice were purchased from Charles River Laboratory (plug date is considered gestational day (GD) 1). The animals were maintained in cages (2-5 per cage) in pathogen-free conditions in the animal facility, with ad libitum access to food and water, and 12L:12D standard conditions. Placentas from both male and female fetuses were used in all experiments and specific effects of fetal sex determined. All the experiments were approved by the Institutional Animal Care and Use Committee at the University of Southern California and CHLA and performed in accordance with the *NIH Guide for the Care and Use of Laboratory Animals*. On gestational day 15, dams were euthanized by CO_2_ inhalation, with death confirmed by cervical dislocation prior to tissue dissection.

### Maternal Immune Activation mouse model and celecoxib treatment

Polyinosinic–polycytidylic acid [poly(I:C)] was prepared by mixing low-molecular-weight poly(I:C) (20 mg/ml; InvivoGen, tlrl-picw) and high-molecular-weight poly(I:C) (1 mg/ml; InvivoGen, tlrl-pic) at a 1:4 ratio (20% LMW, 80% HMW). The resulting mixture was administered to pregnant dams at a final dose of 13.8 mg/kg. Pregnant CD-1 mice were intraperitoneally (i.p.) injected once on GD13 with saline or Poly(I:C). For celecoxib (CX) treatment, pregnant CD-1 mice were (i.p.) injected once on GD 13 with saline or poly(I:C) and subsequently (i.p.) injected with DMSO or celecoxib (20mg/kg) at GD14. Dams were randomly assigned to the following groups: Saline, Saline + CX, pIC, pIC + CX. In these groups, dams were subjected to: 1) live magnetic resonance imaging (MRI) 48 h after injection followed by 2) placenta collection for IHC after MRI scan. A separate cohort of 9 dams was used for placental COX2 activity measures on GD15.

### Magnetic Resonance Imaging

All magnetic resonance imaging (MRI) scans were performed using an MR Solutions 7T PET MR system (bore size ∼24-mm, up to 600 mT.m^−1^ maximum gradient and a 20-mm internal diameter quadrature bird cage mouse head coil. Mice were anesthetized by 1–1.2% isoflurane in air. Respiration rate (80.0 ± 10.0 breaths per min) and body temperature (36.5 ± 0.5 °C) were monitored during the experiments, as we described previously^43^. For abdominal scans, the sequences were collected with respiratory gating in the following order: 1/ Pre- and post-Gadolinium contrast coronal T1-weighted imaging (2D-fast spin echo (FSE), TR/TE (time repetition/time echo) 1,150/11 ms, echo train length 4, 6 averages, 20 slices, slice thickness 400 μm, in-plane resolution 140 x 140 μm^2^) to assess PBB permeability. A bolus dose (140 μL) of 1.5 mmol.kg^−1^ Gd-DTPA (gadolinium diethylenetriamine pentaacetic acid (Gd-DPTA) diluted in saline 1:2) was injected into the tail vein; 2/ T2- weighted imaging (2D-FSE, TR/TE 3,010/45 ms, echo train length 7, 25 slices, slice thickness 1 mm, in-plane resolution 140 x 140 μm^2^) to obtain anatomical images. Total imaging time was approximately 40-50 min per mouse depending upon respiratory gating.

To study PBB permeability, we analyzed the difference in mean signal intensity (SI) between the pre- and post-contrast T1-weighted images using specific regions-of-interest (ROIs) in all studied group^9^. Placentas (which includes both labyrinth and chorionic plate) were manually segmented on high-resolution T2-weighted images using ImageJ.

### COX2 enzymatic activity measurement

Placentas were quickly harvested 28 h after poly(I:C) injection, rinsed in ice-cold phosphate buffer solution (PBS), and frozen in liquid nitrogen. COX2 activity was measured using a COX Activity Assay Kit (Abcam; #ab204699). Samples were processed according to manufacturer instructions; briefly, tissue was transferred to an Eppendorf tube with 200 μL of lysate buffer (protease inhibitor mixture in PBS) and then, homogenized using a plastic tissue homogenizer. COX2 enzymatic activity was shown as μU/mg. Unit definition: 1 Unit COX activity = amount of cyclooxygenase which generates 1.0 μmol of resorufin per minute. at pH 8.0, 25°C. Each sample was assayed in triplicate.

### Placenta homogenates preparation

For western blotting, decidua basalis, junctional zone, and placental labyrinth were carefully dissected on ice from GD15 mouse placentas to isolate maternal and fetal compartments, as previously described (37384519). Briefly, for each mg of tissue, 10 µL of 2x Tris-EDTA buffer (pH 8.3) supplemented with Protease Inhibitor Cocktail (P8340, Sigma-Aldrich) was added, and samples were homogenized on ice for 3 min using a handheld tissue grinder. Further, the same volume of 2x Treatment buffer (125 mM Tris, 4% SDS, 20% glycerol; pH 6.8) was added and samples were heated at 50 °C for 15 minutes and sonicated. Supernatant was collected after centrifugation (20 minutes, 16,000xg, 4 °C). For ELISA measurement of perfused placentas, samples were dissected into decidua basalis and labyrinth, homogenized on ice in 20 µl of 2× Tris-EDTA buffer (50 mM Tris-HCl, 10 mM EDTA, pH 8.3, supplemented with Protease Inhibitor Cocktail (P8340, Sigma-Aldrich)) using a handheld pestle grinder (3 min), and then sonicated. Homogenates were centrifuged at 16,000 g for 20 min at 4 °C, and the supernatant (soluble protein fraction) was collected. Homogenates were stored at -80 °C and protein concentration was determined using Quick Start Bradford Protein Assay Kit 2 (5000202EDU, BioRad).

### Western blot

Homogenates (30µg) from decidua basalis, junctional zone, and labyrinth were mixed with loading buffer under reducing conditions. Samples were heated at 95 °C for 5 min and separated on 4-12% Bis-Tris gel (EP1, EP2, EP3, EP4; NP0321BOX, Thermo Fisher, MES running buffer) or 16% Tricine gel (mPGES1, EC66955BOX, Thermo Fisher, Tricine running buffer) at 130 V. Proteins were transferred to PVDF membranes (iBlot PVDF regular stack, IB401001, 0.2 µm, Thermo Fisher) using iBlot program P0 (7 min). Antibody incubations were performed on the iBind Western device (SLF1000, Thermo Fischer) using either the iBind Fluorescent Detection Solution Kit (mPGES1, EP1, EP2, EP3, EP4; SLF1019, Thermo Fisher) or the iBind Detection Solution Kit (EP3; SLF1020, Thermo Fisher), which provide integrated blocking and wash buffers. Primary antibodies were used: EP3 (rabbit, 1:1000, Cayman Chemical, 101760), mPGES1 (rabbit, 1:200, Cayman Chemical, 160140), b-actin (goat, 1:500 Abcam, ab8229). For confirmation of the EP3 bands, the EP3 antibody was incubated with EP3 receptor blocking peptide (Cayman Chemical, 301760) for 1 hours at RT prior to the primary incubation. IRDye 680RD and 800CW secondaries (Li-COR, anti-rabbit and anti-goat as appropriate, 1:10 000) were used for near-infrared detection on an Odyssey Clx (Li-COR). Blots were normalized to β-actin.

## Dissection and Culture of Placental Labyrinth Explants

Placental labyrinths were dissected as described in the section “Placenta homogenates preparation “. Tissue was cut into 1-1.5 mm^3^ explants, rinsed in cold DMEM/F12, and placed in 12-well plates (10 explants/well) containing 2 mL of DMEM/F12 supplemented with 10% fetal bovine serum, 100 U/mL penicillin and 0.1 mg/mL streptomycin. Explants were incubated at 37 °C under 5% CO_2_ in a sterile incubator for 2 h (recovery phase). Subsequently, explants were cultured for 48 h in either PGE2 (100 pg/mL; Cayman, 14010), the EP3 agonist sulprostone (10 µM; Cayman, 14765), or control medium (DMEM/F12), with media changed after 24 h. In separate experiments, explants were incubated with PGH2 (350 ng/mL; Cayman, 17020), which was added at 0, 12, 24, and 36 h, and explants media were collected at 0, 1, 2, 3, 5, 12, 24, 36 h for analysis.

### Analysis of Placental Explant Viability

Explant viability was assessed at 0, 24 and 48 h using the MTT (thiazolyl blue tetrazolium bromide) assay (Thermo Fisher, M6494). Explants were washed with Opti-MEM and incubated in 500 µg/ml MTT solution at 37 °C for 2 h. Following incubation, 500 µL of DMSO was added per well and plates were shaken for 15 minutes at room temperature. Absorbance was measured at 540 nm and normalized to explant tissue weight (mg). Explants integrity was further evaluated by LDH assay kit (Cayman, 601170) according to the manufacturer’s instructions. The results are presented as percentage of LDH release relative to positive control. Data are shown in Supplementary Fig. 1.

### *Ex vivo* placenta perfusion

One placenta per dam was collected at GD14 and transferred immediately to a thermostatically controlled incubation chamber at 37°C. The uterine artery was cannulated with a 200-mm ID catheter and perfused at 20 mL/min with DMEM (Life Technologies, 11054-001) in the presence or absence of arachidonic acid (2.5 mg/mL; Cayman, 90010). The umbilical artery was cannulated with a 105-mm ID catheter and perfused at 5 mL/min with DMEM. Perfusates were collected from the umbilical vein over 90 min and analyzed for PGH2 or PGE2 concentration by ELISA. Detailed methodology for *ex vivo* placenta perfusion has been previously described^15^.

### PGE2 and PGH2 ELISA

Fetal perfusates from *ex vivo* placenta perfusions and culture media from untreated explants or explants treated with PGH2 were collected, and immediately frozen at -80°C until analysis. PGE2 concentrations were measured using an ELISA kit (ENZO Life Science, ADI-930-001) according to manufacturer’s instructions. PGH2 concentrations were measured both in fetal perfusates from *ex vivo* placenta perfusions and in placental tissue harvested post-perfusion. PGH2 concentrations were measured using a PGH2 ELISA kit (assay sensitivity: 0.97 pg/mL; lower detection range: 3.13 pg/mL; AFG Bioscience, EK24759) according to the manufacturer’s protocol.

### Immunohistochemistry

Mouse placentas and placental explants were fixed in 4% PFA in PBS overnight at 4°C, cryoprotected in 10%, 20% and 30% sucrose (24 h each), embedded in OCT, sectioned at 20 μm with a cryostat (CM3050S, Leica), mounted on Superfrost Plus slides, and stored at - 80°C. Human placenta samples were paraffin-embedded and cut into 5 μm sections; slides were deparaffinized in xylene (20 min), and rehydrated through graded ethanol (100%, 95%, 70% and 50%) for 10 min each. Antigen retrieval was performed in 10 mM sodium citrate buffer (pH 6.0) at 95 °C for 10 min. For IHC, mouse, human, and explant sections were permeabilized and blocked in PBS with 0.1% Triton X-100 and 2%-5% fetal bovine serum (FBS) for 2 h at room temperature, incubated overnight at 4°C with primary antibodies, and then incubated with secondary antibodies for 2 h at room temperature. Primary antibodies included CD31 (PECAM-1, hamster, 1:1000, Millipore, MAB1398Z), CD31 (Pecam-1, mouse, 1:500, Abcam, Ab9498), CD31 (Pecam-1, rat, 1:100, BD Biosciences, 550274), anti-alpha smooth muscle actin (SMA; mouse, 1:100, Dako, M0851 or rabbit, 1:1000, Abcam, AB124964 or rabbit monoclonal, 1:500, Abcam, Ab32575), CD68 (rat, 1:200, Bio-Rad, MCA1957GA), CD45 (rat, 1:50, BD Biosciences, 550539), CD206-Biotin (1:200, Bio-Rad, MAC2235B), COX2 (rabbit, 1:200, Cayman, 160126), MCT1 (chicken, 1:1000, Millipore, AB1286-I), mPGES1 (rabbit, 1:200, Cayman Chemical, 160140), EP3 (rabbit, 1:200, Cayman Chemical, 101740). Secondary antibodies raised in donkey (Jackson ImmunoResearch) were used at 1:800, including Alexa 488 anti-rabbit/hamster/rat, Alexa 647 anti-rat and Rhodamine Red anti-chicken/hamster/mouse, as well as HRP-conjugated anti-mouse, biotin-conjugated anti-rabbit, and DyLight 549 streptavidin (1:500, Vector SA-5549). For SMA, an additional incubation with Tyramide Signal Amplification-Cy3 (1:100, PerkinElmer NEL7441B001KT) was performed for 30 min.

### Image acquisition, quantification and analysis

Sections were coverslipped with Prolong Gold (Vector) and imaged using a Zeiss Axioimager 2 microscope or Zeiss LSM800 confocal microscope, using identical acquisition settings within staining for group comparisons.

In each mouse, three regions of the decidua basalis or labyrinth (left, middle and right) were analyzed in four non-adjacent sections (∼100 µm apart) and averaged per animal. Quantification of pericyte-like cell coverage of endothelial cell was performed in ImageJ as described previously^9^. Briefly, SMA/CD31 32-bit images were thresholded using the Otsu plugin, and the ratio of SMA to CD31 area was calculated. For macrophage assessment, CD45 and CD68/CD206 double-positive cells were counted manually in defined ROIs using Leica Microscope Imaging Software (REF) and normalized per mm^2^. In mouse labyrinth explants, six ROIs from each explant in non-adjacent sections (∼100 µm apart). Explants were prepared from multiple placentas within a litter, and results were averaged so that each litter was considered one biological replicate. Data were obtained from at least three independent litters. Pericyte coverage was quantified as above. For human placentas, Akoya multiplex immunofluorescence assay with AI-based signal quantification was used.

### Human placenta samples collection

Histological samples of placentas at 33-41 weeks, from patients asymptomatic for SARS-CoV-2 (8 patients), patients with chronic and antepartum infections (6 patients) and patients without infection (5 patients) were collected from Los Angeles General Medical Center. ‘Other infections’ category included HIV, HSV, chronic hepatitis B, prior syphilis, and chorioamnionitis/chorionitis. Patients with COVID-19 had a positive RT-PCR test for SARS-CoV-2 prior to delivery. Matched controls had negative testing for SARS-CoV-2 and maternal HIV and were without clinical history of perinatal infections. Maternal clinical data were blinded to the investigators. Placental specimens were fixed with 4% formaldehyde, embedded in paraffin and sectioned (5 mm) onto slides at the USC Pathology Core. All participating women provided written informed consent prior to sample collection (USC IRB approval number HS-20-00359). The demographic and clinical characteristics of the study groups are shown in Supplementary Table 1.

### Multiplex Immunofluorescence (mIF) assays

A six-plex mIF assay consisting of CD3, CD31, SMA, COX2, CD68 and DAPI was developed for analysis. Protocol development and subsequent analysis were based on the technique previously reported by Taube et al^44^. In brief, the staining protocol was applied to three placenta cases, and each marker was analyzed for target range of 10–30 in normalized brightness counts based on the Akoya Biosciences Phenochart Intensity Cursor function. The concentration of each antibody was adjusted until brightness intensity counts were satisfactory for all markers. The pairing of antibody with fluorophore and order of staining was based on the principal of pairing the strongest antibody expresser with the weakest expressing fluorophore (brightness) and vice versa. Each antibody was tested in the 1st, 3rd, and last position to evaluate the signal strength of all markers across various position in the multiplex panel. The final protocol order used to stain the tissues is provided in the table below. Following panel optimization, the cases were stained according to protocol.

| Order | Antibody (clone) | 1°DF | Opals | Opal DF |
| --- | --- | --- | --- | --- |
| 1 | CD3 (LN10) | RTU* | Opal 480 | 1:100 |
| 2 | CD31 (JC70A) | RTU* | Opal 520 | 1:100 |
| 3 | SMA (1A4) | 1:500 | Opal 690 | 1:100 |
| 4 | COX2 (ABCAM 15191) | 1:100 | Opal 570 | 1:100 |
| 5 | CD68 (KP1) | 1:1000 | TSA-DIG Opal 780 | 1:50, 1:25 |

\*RTU = Ready to use

The standard six-color TSA protocol template on the Leica BOND RX was used. All tissue sections underwent an initial 1-hour baking step at 65°C. A second bake and de-wax step was then performed using a dewax solution (AR9222, Leica Biosystems) on the BOND RX to ensure that all paraffin was removed. Slides were then stained using the automated mIF staining protocol.

### mIF image analysis

Multispectral images of the stained sections were acquired using the Akoya Biosciences PhenoImager Automated Quantitative Pathology Imaging System. Prior to processing, all images were assessed for quality control. All images were processed and analyzed with inForm software (V.2.6.10). A single algorithm for spectral unmixing, cell segmentation, cell classification, phenotyping, and quantification of expression intensity for each marker was developed. Cells were segmented into cytoplasmic, nuclear, and membrane compartments. All images were analyzed using the Batch Process protocol in inForm and exported for analysis in the R-script package phenoptrReports (Akoya BioSciences). The phenoptyrReports analysis was used to provide information on density, location, intensity, co-localization, and spatial relationships for all biomarkers.

## Statistical Analysis

Statistical analyses were performed using GraphPad Prism 8.0 software (La Jolla, CA). All data were tested for normality test. Results from histological and MRI experiments were compared using unpaired two-tailed t-test, one-way ANOVA followed by Tukey’s post-hoc comparison analysis, or two-way ANOVA followed by Bonferroni’s correction, depending on the number of variables included in the comparison. mPGES1 protein expression is evaluated by simple t-test. EP3 protein expression and explants data were evaluated by unpaired two-tailed t-test. Human data were evaluated by Kruskal-Wallis test with Dunn’s multiple comparison and presented as median + IQR. All other data are presented as mean ± SEM. Test type and levels of statistical significance are indicated in the figure legends.

