## Supplemental materials for "Placental prostaglandin signaling disrupts barrier integrity and relays an acute inflammatory signal to the fetus"

### Supplementary data

**Supplementary table 1 | Clinical data of human placenta samples.**

| | Slide number | Gestational age (weeks) | Infection type |
|---|---|---|---|
| **Controls** | | | None |
| | 1286 | 40 | None |
| | 1290 | 40 | None |
| | 1339 | 37 | None |
| | 2237 | 41 | None |
| | 2713 | 35 | None |
| **Cases** | 2324 | 34 | HIV- undetectable HSV viral load |
| | 2407 | 38 | Chorioamnionitis |
| | 2270 | 38 | HIV - undetectable viral load. Chronic Hepatitis B |
| | 0040 | 37 | Chorionitis |
| | 2214 | 37 | HIV – unknown viral load |
| | 2251 | 36 | HIV, HSV |
| **COVID-19** | 2736 | 33 | Covid - asymptomatic |
| | 2704 | 38 | Covid - asymptomatic |
| | 2771 | Unknown | Covid - asymptomatic |
| | 2787 | 35 | Covid - asymptomatic |
| | 2899 | 34 | Covid - asymptomatic |
| | 3392 | 39 | Covid - asymptomatic |
| | 2558 | 39 | Covid - asymptomatic |
| | 2272 | 39 | Covid - asymptomatic |

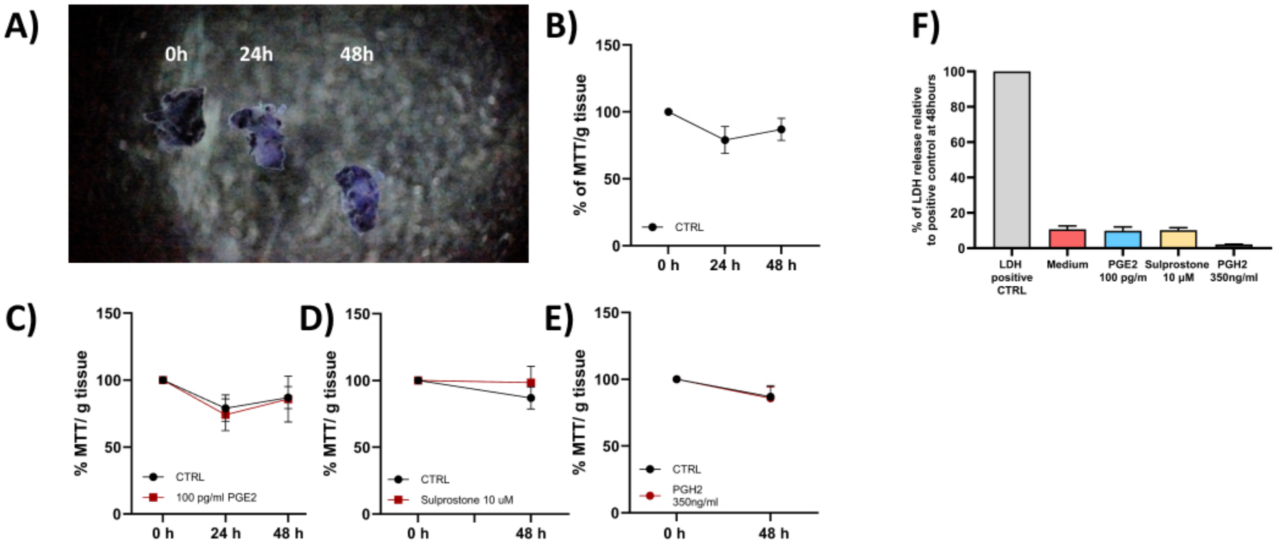


**Supplementary figure 1 | Viability and integrity assessment of *ex vivo* explants culture.** Representative picture of placental labyrinth explants after incubation with MTT+ (A) Quantification of MTT+ assay in non-treated explants (B), PGE2 (100 pg/ml) treated explants (C), Sulprostone 10 $\mu$M treated explants (D) and PGH2 (350 ng/ml) treated explants (E). Quantification of LDH assay expressed as a percentage of LDH release relative to positive control at 48 hours (F). Data are presented as mean + SEM. n=4.

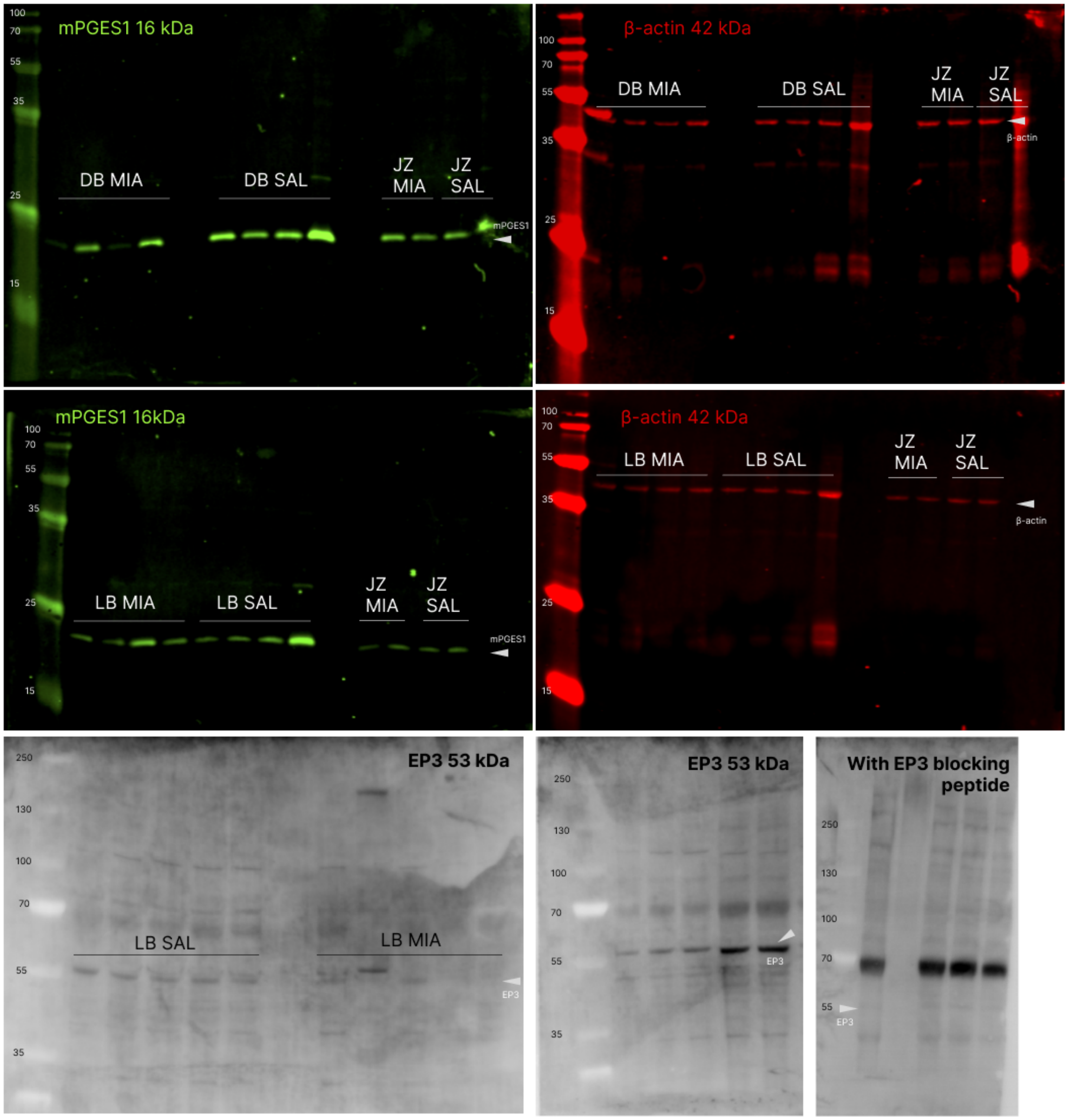


**Supplementary figure 2 | Uncropped immunoblots of proteins analyzed (including mPGES1, β-actin, EP3)**

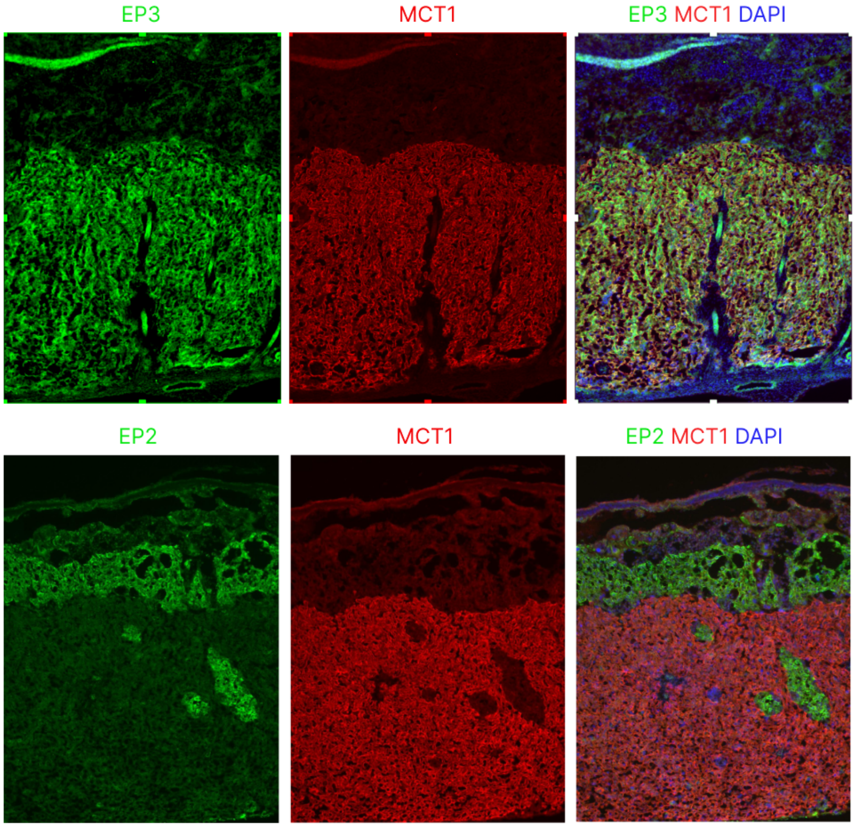


**Supplementary figure 3 l Representative images showing EP3 (green) overlap with MCT1 (red) and EP2 (green) absence of overlap with MCT1 (red).**
